# Exposure of zebra mussels to extracorporeal shock waves demonstrates formation of new mineralized tissue inside and outside the focus zone

**DOI:** 10.1101/254771

**Authors:** Katharina Sternecker, Juergen Geist, Sebastian Beggel, Kristin Dietz-Laursonn, Matias de la Fuente, Hans-Georg Frank, John P. Furia, Stefan Milz, Christoph Schmitz

## Abstract

A substantial body of evidence supports the use of extracorporeal shock wave therapy (ESWT) for fracture nonunions in human medicine. However, the success rate (i.e., radiographic union at six months after ESWT) is only approximately 75%. Detailed knowledge regarding the underlying mechanisms that induce bio-calcification after ESWT is limited. The aim of the present study was to analyze the biological response within mineralized tissue of a new invertebrate model organism, the zebra mussel *Dreissena polymorpha*, after exposure with extracorporeal shock waves (ESWs). Mussels were exposed to ESWs with positive energy density of 0.4 mJ/mm2 or were sham exposed. Detection of newly calcified tissue was performed by concomitantly exposing the mussels to fluorescent markers. Two weeks later, the fluorescence signal intensity of the valves was measured. Mussels exposed to ESWs showed a statistically significantly higher mean fluorescence signal intensity within the shell zone than mussels that were sham exposed. Additional acoustic measurements revealed that the increased mean fluorescence signal intensity within the shell of those mussels that were exposed to ESWs was independent of the size and position of the focal point of the ESWs. These data demonstrate that induction of bio-calcification after ESWT may not be restricted to the region of direct energy transfer of ESWs into calcified tissue. The results of the present study are of relevance for better understanding of the molecular and cellular mechanisms that induce formation of new mineralized tissue after ESWT. Specifically, bio-calcification following ESWT may extend beyond the direct area of treatment.

**Summary statement:** The use of zebra mussels in research on extracorporeal shock wave (ESW) therapy for fracture nonunions allows new insights into the complex process of induction of biomineralization by ESWs.

## Introduction

Extracorporeal shock wave therapy (ESWT) is an accepted treatment for fracture nonunions (reviewed in, e.g., Haupt, 1997; Kertzman et al., 2017; Schaden et al., 2015). Overall, the mean success rate reported in these studies (defined as radiographic union at six months after ESWT) was only approximately 75%. There is no generally accepted protocol nor is there a consensus as to which method of extracorporeal shock wave (ESW) production, focal or radial, is optimal. Hence, better understanding of the molecular and cellular mechanisms of action of ESWs on mineralized tissues is paramount.

These mechanisms are only partially understood (Chamberlain and Colborne 2016; Cheng and Wang 2015; Zelle et al., 2010). Early studies focusing on ESW-induced micro-cracks of bone (i.e., cracks that can only be detected under a microscope) indicated that ESWs can induce reorganization and regeneration of bone via callous production and subsequent secondary bone healing (Da Costa Gómez et al., 2004; Ikeda et al., 1999; Valchanou and Michailov, 1991). Further, Tischer et al. (2008) demonstrated that ESWs can also induce new bone formation without generation of micro-cracks that is comparable to primary bone healing.

ESWT is generally considered to be a technically simple procedure. The goal is to position the focal zone, the area of greatest wave concentration, in the area of the nonunion. One potential technical error when using this procedure is the improper positioning of the focus zone thereby missing the fracture gap. On the other hand, Tischer et al. (2002) demonstrated on rabbits that ESWs can induce new bone formation inside and outside the focus zone. In fact, the important question about correlations between the regional, three-dimensional (3D) distribution of pressure generated by ESWs and the regional (3D) pattern of new bone formation induced by ESWs could not be answered so far. This is due to the fact that for technical reasons, it is not possible to measure the regional (3D) distribution of pressure generated by ESWs within a limb of a human or one of the commonly used vertebrate animal models in ESWT research (goat, rabbit, rat and mouse) *in vivo*.

The aim of the present study was to test the hypothesis that exposure of invertebrate mussels to ESWs results in formation of new mineralized tissue in the mussel shell, with a regional (3D) pattern of new mineralized tissue that correlates to the regional (3D) distribution of pressure generated by the ESWs (i.e., higher amounts of new mineralized tissue at regions of higher pressure). Such a finding would suggest that during ESWT, the operator of the ESWT device has to position the focus point most exactly to cause wanted positive results.

## Materials and methods

### Animals

The present study was performed on zebra mussels (*Dreissena polymorpha*). *Dreissena polymorpha* has been identified as a promising model organism for studying biomineralization e.g. in ecotoxicology (Immel et al., 2016). *Dreissena polymorpha* typically occurs in high population densities, has been globally introduced, can genetically be unambiguously identified (Beggel et al., 2015) and is an invertebrate facilitating its unlimited use in animal experiments compared to vertebrates. All of these aspects render *Dreissena polymorpha* a readily available and ideal target organism.

Forty-eight mussels were used for investigating the formation of new mineralized tissue after exposure to ESWs. These 48 mussels were collected by hand from the river Ischler Ache (Upper Danube Drainage, Bavaria, Germany) in July 2014, i.e., before spawning and the peak of the natural growth season in late summer (Jantz and Neumann, 1998). Mussels were individually housed in separate chambers of six-well multiwell plates at the Aquatic System Biology Unit, Technical University of Munich (Freising, Germany). Two multiwell plates each were fixed in a bucket filled up with 10 l ground water (mean temperature = 12.5° C). Acclimatization to experimental temperature conditions was performed according to ASTM E2455-06 (Ingersoll et al., 2006) by gradually decreasing the temperature by no more than about 3° C per hour. Approximately 30% of the water in the buckets was changed daily. Mussels were fed by adding 0.2 ml/l Shellfish Diet 1800 (Reed Mariculture, Campbell, CA, USA) to the incubation water once per week.

Two additional mussels were used for characterizing zebra mussel morphology and for acoustic measurements. These mussels were collected by hand from the Lake Ammer (Bavaria, Germany) in July 2015 and the Lake Starnberg (Bavaria, Germany) in April 2016.

Mussels were genetically validated to be *Dreissena polymorpha* following the restriction fragment length polymorphism (RFLP) method described by Beggel et al. (2015).

All experiments were performed according to German animal protection regulations which do not require registration or approval of experiments with zebra mussels.

### Experiments performed on the mussels used for investigating the formation of new mineralized tissue after exposure to ESWs

The 48 mussels used for investigating the formation of new mineralized tissue after exposure to ESWs were randomly divided into four groups (A to D; n=12 mussels per group). Two weeks before exposure to ESWs or sham exposure the size dimension of each mussel shell were determined using digital photography and quantitative analysis with the software AxioVs40 (version 4.8.2.0; Zeiss, Jena, Germany). Specifically, length, height and width of the mussel shells were measured to the nearest 0.1 mm as described by Claxton et al. (1997) (Fig. 1A,B). The mean shell length was 22.7 ± 0.3 mm (mean ± standard error of the mean; SEM), the mean shell width was 11.2 ± 0.1 mm and the mean shell height was 11.8 ± 0.2 mm. No statistically significant differences between the groups were observed (Kruskal-Wallis test; p>0.05).

**Fig. 1.**
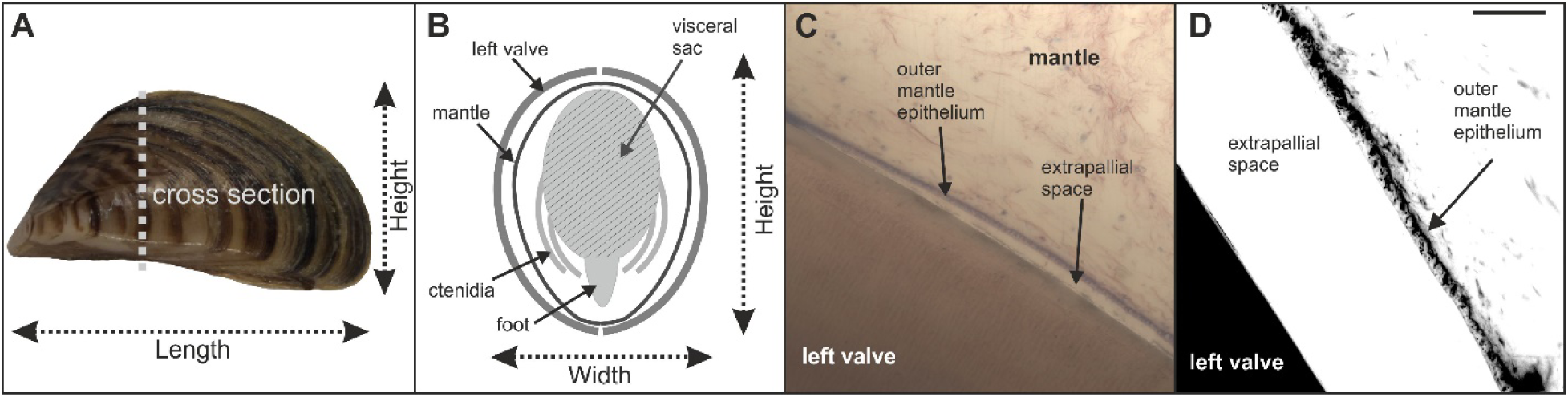
Morphology of the zebra mussel *Dreissena polymorpha*. (A) Side view on a zebra mussel. (B) Schematic of a cross section through a zebra mussel. The soft tissue (visceral sac with internal organs, foot and ctenidia (i.e. gills)), mantle and shell are indicated. (C, D) The biocalcification zone of a zebra mussel imaged with brightfield microscopy (C) and fluorescence microscopy (D). Details are provided in the main text. The scale bar in (D) represents 5 mm in (A), 100 µm in (C) and 25 µm in (D).

For histologic detection of modifications in the mussel shell after exposure to ESWs or sham exposure, mussels in Groups A and B were exposed to the fluorescent calcium binding dyes xylenol orange and calcein (O’Brien et al., 2002; Pautke et al., 2005; Rahn and Perren, 1971; Suzuki and Mathews, 1966). Fluorescent calcium binding dyes allow an easy identification of newly formed mineral deposits and were previously used in various *in vivo* studies on the formation of new bone after exposure of laboratory animals to ESWs (Delius et al., 1995; Gollwitzer et al., 2013; Tischer et al., 2008). The excitation wavelengths of xylenol orange are 440 and 570 nm, and its emission wave length is 610 nm. For calcein the corresponding values are 494 nm (excitation wavelength) and 517 nm (emission wavelength) (Pautke et al., 2005).

Mussels in Groups A and B were incubated in xylenol orange (90 mg/l; Sigma-Aldrich, St. Louis, MO, USA) for 24 h, shortly after determining the dimension of the mussel shells. Two weeks later, mussels in Group A were exposed in a customized 40 l water tank filled up with ground water to 1000 ESWs generated with an ESWT device (Swiss Piezoclast; Electro Medical Systems S.A., Nyon, Switzerland) and piezoelectric ESW applicator (F10G4; Richard Wolf, Knittlingen, Germany). The positive energy density (ED_+_) of the ESWs was 0.4 mJ/mm^2^ and the pulse rate of the ESWs was 8 Hz. Extracorporeal shock waves were applied on the left valve of the mussels. This was achieved by positioning the mussels in a custom-made nylon fine-mesh net (<1 mm mesh size) at a distance of 45 mm to the applicator. Using these settings the midpoint of the left valve of the mussels was placed at the position of the focus point of the ESWs, i.e., at the position of maximum energy (Fig. 2). Immediately after exposure to ESWs the mussels were incubated in calcein (10 mg/l; Sigma-Aldrich, St. Louis, MO, USA) for 24 h.

**Fig. 2.**
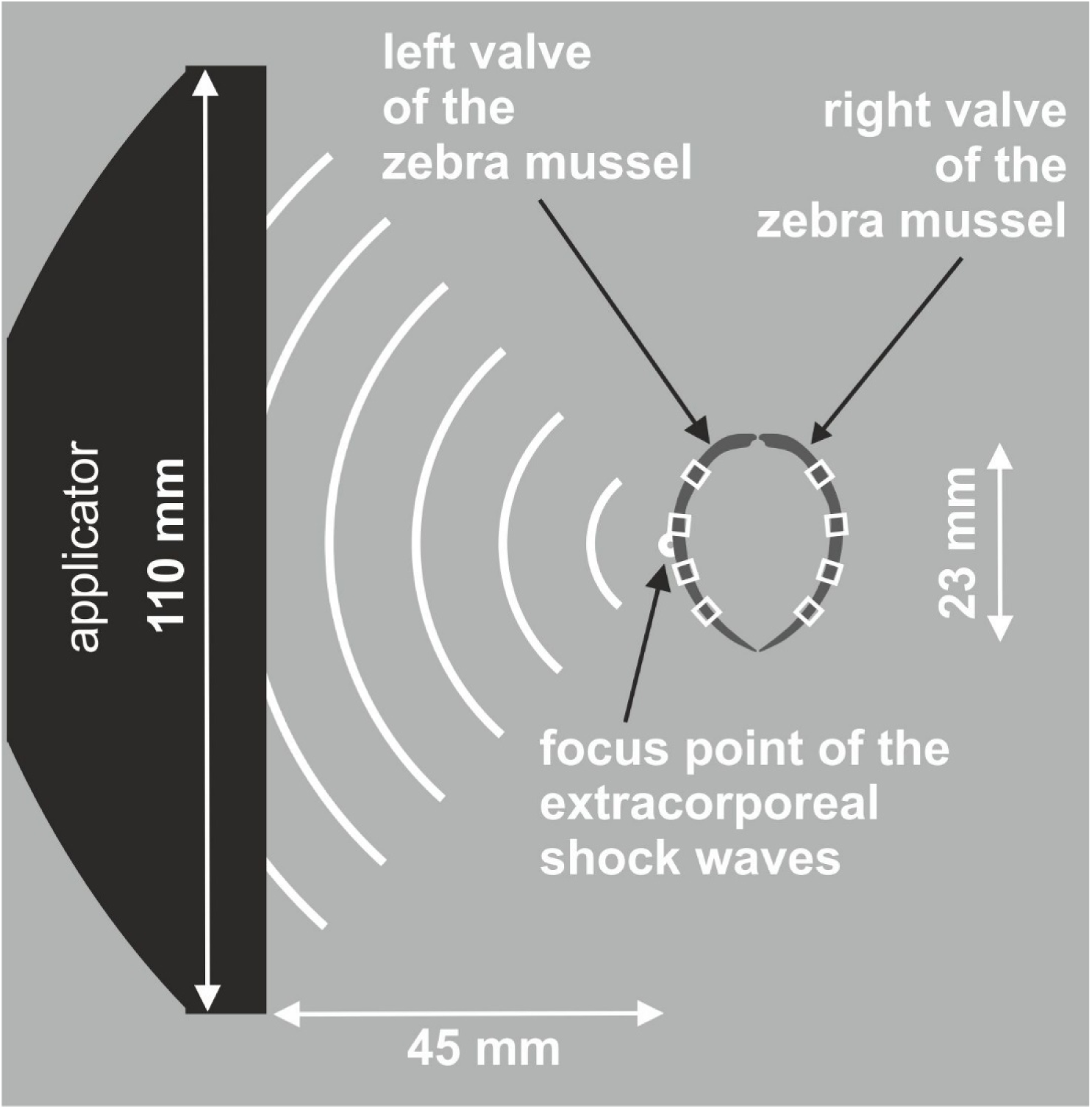
Principle of the experimental setup used for exposing zebra mussels to extracorporeal shock waves (ESWs). The distance between the focus point of the ESWs and the applicator of the ESWT device was determined using shadowgraph imaging as described in detail in Schmitz et al. (2013). Then the left valve of each mussel was placed at the focus point using a custom-made fine nylon mesh net (not shown). The white rectangles over the left and right valves indicate the positions on the mussel shell where formation of new mineralized tissue after exposure to ESWs or sham exposure was investigated.

During fluorescent marker incubation the water volume in the buckets was reduced to 2 l in order to minimize toxic waste caused by the dye solutions, and water was completely changed after 24 h incubation.

Mussels in Group B were exactly treated like mussels in Group A but were sham exposed to ESWs. This was achieved by not switching on the ESWT device.

Mussels in Groups C and D were exposed to ESWs (Group C) or sham exposed (Group D) as described above, but not incubated in xylenol orange and calcein.

Formation of new mineralized tissue after exposure to ESWs or sham exposure was investigated on the mussels in Groups A and B. Mussels in Groups C and D were used to determine mortality rates after exposure to ESWs or sham exposure, with zero mortality observed in these groups over two weeks. The mortality in Group B (incubation in fluorescent dyes; sham exposure to ESWs) was also zero during the time period of the experiments. In Group A (incubation in fluorescent dyes; exposure to ESWs) no mussel died before or immediately after exposure to ESWs. On the other hand, two mussels in Group A died during the two-week period following exposure to ESWs. Nevertheless, it was possible to analyze the valves of these mussels as the valves of the other mussels in Groups A and B.

### Histologic processing and analysis of mussels used for investigating the formation of new mineralized tissue after exposure to ESWs or sham exposure

Two weeks after exposure to ESWs or sham exposure mussels in Groups A and B were sacrificed by immersion fixation in 70% ethanol for 48 h. Mussel shell (left and right valves) and soft tissue were separated and mussel valves were dehydrated in ascending ethanol fractions (70% for three days, 80% for six days, 90% for eight days and 100% for 14 days). After defatting with xylene for one week and incubation in methanol for two weeks, both valves were embedded in methyl methacrylate (Sigma-Aldrich, Buchs, Switzerland) according to Milz and Putz (1994). After curing, serial, transverse 400-µm thick sections of the valves were cut across the longest growth axis (Fig. 3A,B) using a saw microtome (SP 1600; Leica, Wetzlar, Germany), yielding the highest resultion of microgrowth patterns (Geist et al., 2005). Shell sections were subsequently ground and polished using a 400 CS micro grinder (EXAKT Advanced Technologies, Norderstedt, Germany). The mean final section thickness was 269 ± 11.5 µm (mean ± SEM), determined in the middle of the sections using a Digimatic micrometer (Mitutoyo, Kawasaki, Japan).

**Fig. 3.**
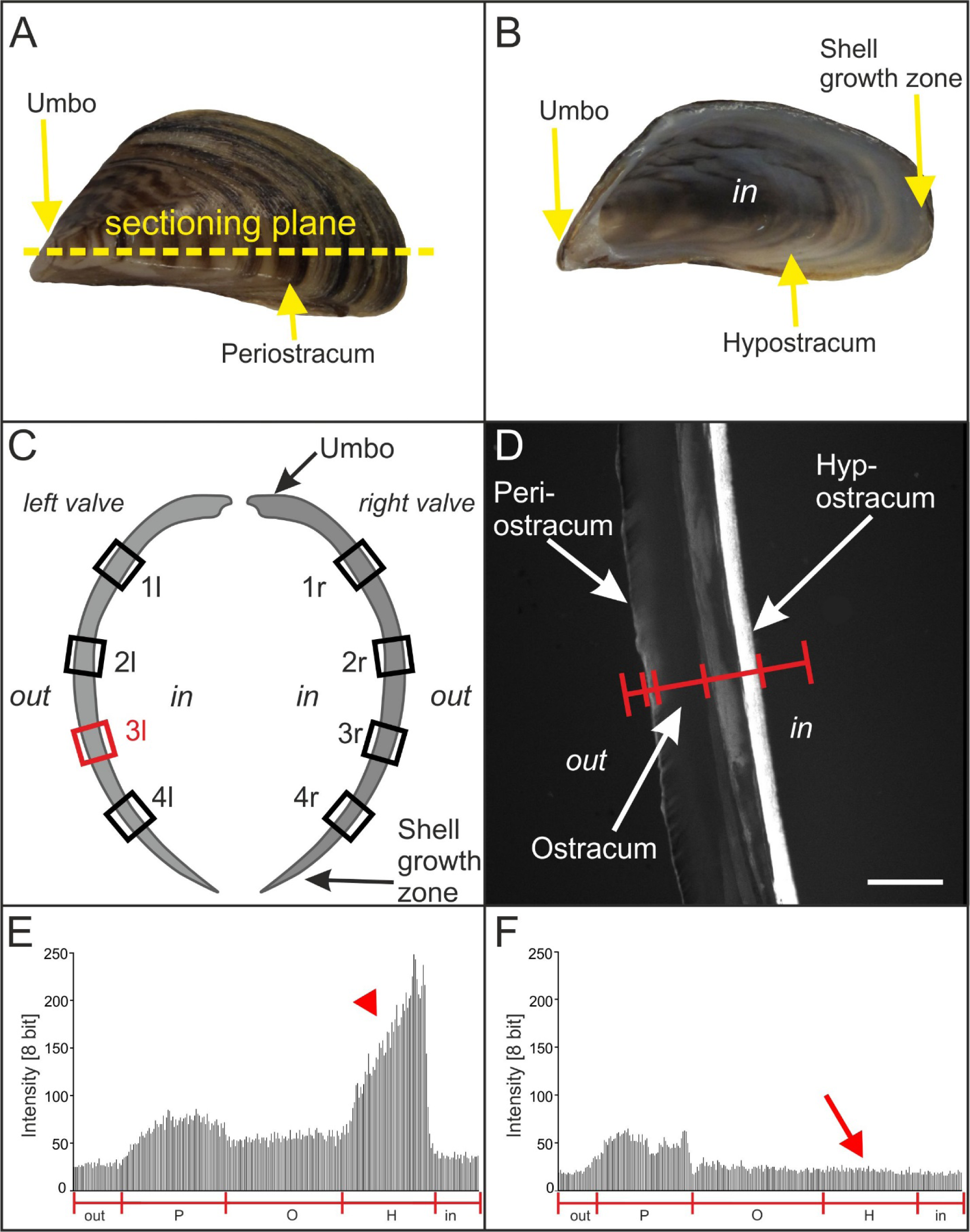
Principle of analyzing the effects of extracorporeal shock waves on zebra mussels. (A) Outer surface of the left valve of a zebra mussel. The umbo, periostracum and the sectioning plane used in the present study are indicated. (B) Inner surface of the left valve of a zebra mussel. The umbo, hypostracum and shell growth zone are indicated. (C) Sketch of the left and right valve of a zebra mussel, with the sites indicated where formation of new mineralized tissue after exposure to ESWs or sham exposure was investigated. The red rectangle (3l) indicates the position at which the measurements shown in (E, F) were performed. (D) Principle of investigating the formation of new mineralized tissue after exposure to ESWs or sham exposure using fluorescence microscopy by determining the fluorescence signal intensity (calcein fluorescence imaging) along a line spanning the entire thickness of the mussel valve. The periostracum, ostracum and hypostracum are indicated. (E) Representative linear pixel plot of the fluorescence signal intensity at position 3l shown in (C) along the red line shown in (D) on the left valve of a mussel exposed to ESWs (Group A) (out, region outside the mussel; P, periostracum, O, ostracum; H, hypostracum; in, region within the mussel). Note the high fluorescence signal intensity at the position of the hypostracum (red arrowhead). (F) Representative linear pixel plot of the fluorescence signal intensity at position 3l shown in (C) along the red line shown in (D) on the left valve of a mussel that was sham exposed (Group B). Note the lack of fluorescence signal intensity at the position of the hypostracum compared to (E) (red arrow). The scale bar in (D) represents 5 mm in (A, B) and 280 µm in (D).

One transverse section of the left and the right valve of each mussel in Groups A and B was investigated with a fluorescence microscope (Olympus BX51WI; Olympus, Tokyo, Japan) using an UPLSAPO4X objective (4×; numerical aperture [N.A.] = 0.16) (Olympus), Alexa Fluor 488 filter (excitation: 498 nm, emission: 520 nm, calcein fluorescence imaging; Chroma, Bellows Falls, VT, USA) as well as Alexa Fluor 594 filter (excitation: 590 nm, emission: 617 nm; xylenol orange fluorescence imaging; Chroma), a gray scale EM CCD camera (model C9100-02, 1000 x 1000 pixels, Hamamatsu Photonics, Hamamatsu City, Japan) and a SOLA LED lamp (Lumencor, Beaverton, OR, USA). On each section the fluorescence signal intensity was measured at four positions (Fig. 3C): next to the umbo (named “1 left” [1l] and “1 right” [1r]), next to the shell growth zone (“4l” and “4r”), and two positions in between (“2l” and “2r” as well as “3l” and “3r”). The umbo itself was excluded from the analysis because substantial autofluorescence was found in this zone caused by the ligament (Fig. 4) (the ligament at the umbo opens the mussel shell and hence, is the antagonist of the adductor muscles that closes the mussel shell).

**Fig. 4.**
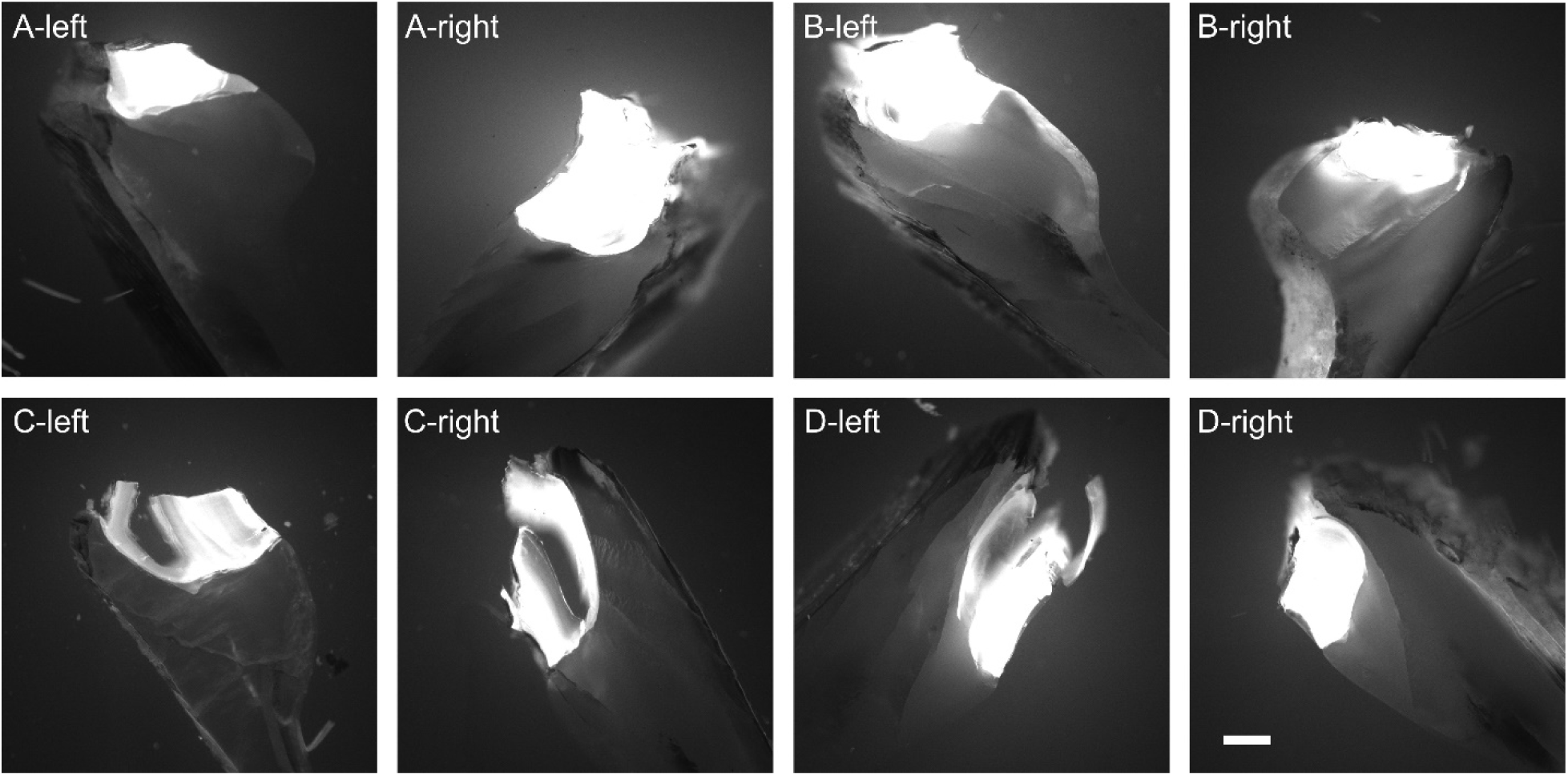
Autofluorescence at the umbo of zebra mussels. The panels show representative photomicrographs (calcein fluorescence imaging) at the umbo of sections of the left and right valve of mussels in Groups A to D. Note that mussels in Groups A and B were incubated in xylenol orange for 24 h and two weeks later in calcein for 24 h. In contrast, mussels in Groups C and D were not incubated in xylenol orange and calcein. The scale bar in the lower right panel represents 250 µm in all panels.

At each position 1l to 4l and 1r to 4r shown in Fig. 3C measurements of fluorescence signal intensity were performed using the Linear Pixel Plot function of the software Stereo Investigator (version 11.07; MBF Bioscience, Williston, VT, USA) along a line spanning the entire mussel valve (Fig. 3D). The outermost layer of the mussel shell is the periostracum (indicated in Fig. 3D) being an organic layer that protects the mussel shell against environmental impact, e.g. acidic pH. The periostracum has an important role during new shell formation at the shell growth zone, but is not involved in building the new hypostracum layers (e.g. Checa, 2000; Petit et al., 1980). The latter also applies to the ostracum (also indicated in Fig. 3D), which is characterized by an inorganic mineralized matrix (aragonite) (Immel et al., 2016). Therefore these layers were excluded from the quantitative analysis of the linear pixel plots. The hypostracum is the nacreous layer of the mussel shell growing by apposition of nacre and showed the highest fluorescence signal intensity in the experiments described in the present study (Fig. 3D). During the process of growing by apposition of nacre organic polymers give structural support, thereby having similar function in starting, organizing and limiting calcification as the organic matrix collagen in vertebrate bones (Beedham, 1964; Mann, 1988). The hypostracum was investigated in the quantitative analysis of the linear pixel plots.

For calcein fluorescence imaging the camera was calibrated by imaging a mussel valve showing high fluorescence signal intensity (taken from Group A; highest fluorescence signal intensity found at the hypostracum). The camera was adjusted so that all pixels of the linear pixel plot (spanning the entire mussel valve) showed an intensity of less than 255 (maximum at 8 bit), resulting in the following configuration: exposure time = 20 ms, sensitivity = 80 and gamma = 1.0. This configuration was kept constant in all subsequent imaging experiments.

Representative linear pixel plots at positions 1l of one mussel each in Groups A and B are shown in Fig. 3E, F. Because of the fact that at positions 1 more pixels represented the hypostracum than at positions 4, individual average fluorescence intensity values were calculated for each position 1 to 4 of each investigated valve. Fluorescence intensity values are presented as dimensionless variable.

### Histologic processing and imaging of a mussel for demonstrating zebra mussel morphology

One muscle was sacrificed, embedded and cut into sections (including grinding and polishing of the sections) as described above for the mussels used for investigating the formation of new mineralized tissue after exposure to ESWs or sham exposure. However, in this case, soft and hard tissue were not separated before embedding. Sections were stained with Giemsa staining. Imaging of the sections was performed with a BX51WI microscope (Olympus) operated in brightfield mode, equipped with an UPLSAPO20X objective (20×, N.A. = 0.75) (Olympus) and Retiga 2000R CCD camera (Q-Imaging, Surrey, BC, Canada).

### Histologic processing of a mussel for acoustic measurements

As the setup of *in vitro* experiments on ESWT can significantly influence the pressure field (Dietz-Laursonn et al., 2016), acoustic measurements were performed using another mussel. The shell length, width and height (24 × 13 × 12 mm) of this mussel were similar to the corresponding mean data of the mussels used for investigating the formation of new mineralized tissue after exposure to ESWs or sham exposure. This mussel was sacrificed by cutting the ligament and separating the soft tissue from the shell. After decontamination in 70% ethanol for 30 min the mussel shell was stored in PBS buffer until acoustic measurements were performed. This was done in order to retain the structural integrity of the mussel shell after death. Thus, the acoustic measurements were conducted most similarly to the experiments performed for investigating the formation of new mineralized tissue after exposure to ESWs or sham exposure.

### Acoustic measurements

Acoustic measurements were carried out at the laboratory of the Chair of Medical Engineering, RWTH Aachen University (Aachen, Germany) according to IEC-61846:1998 (Ultrasonics - Pressure pulse lithotripters - Characteristics of fields) in a cylindrical water tank (diameter = 50 cm) using a fiber optic probe hydrophone (FOPH 2000; RP acoustics, Leutenbach, Germany) with a bandwidth of 100 MHz coupled to an oscilloscope (DPO 2024; Tektronix, Beaverton, OR, USA). Positioning of the FOPH was controlled with a XYZ-positioning table, allowing a resolution of the position of 12.5 µm. During all acoustic measurements the Z-axis was parallel to the acoustic axis. Measurements were carried out with the same piezoelectric ESWT device and a different but technically identical ESW applicator as used in the biologic experiments.

Pressure was calculated from the voltage that was recorded by the oscilloscope according to the specifications of the manufacturer of the hydrophone. The energy density (ED) of the ESWs was calculated from the pressure as

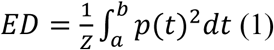

with *Z* the impedance of sound in water (1.5×10^6^ kg m^−2^s^−1^), *p*(*t*) the pressure as a function of time and the integration limits *a* and *b*. The limits of the positive ED (ED_+_) were defined according to IEC-61846:1998. The negative ED (ED_−_) was calculated accordingly for the negative pressure part.

A first series of acoustic measurements (depicted in Fig. 5A) was dedicated to determining the position of the focus point within the 3D pressure field generated by the ESWs (indicated as “Pos1” in Fig. 5) (note that Pos1 approximately corresponded to the position of the middle of the left valve of a mussel in case the entire mussel shell were positioned in the 3D pressure field). To this end, the water tank was filled with tap water (mean temperature during the measurements: 11.5 ± 1.0 °C; temperature controlled with a thermostat, ice and a pond pump; water level 14.5 cm above the baseplate of the water tank, i.e. 10 cm above the focus point). To take into account possible influences of the experimental setup to the pressure distribution, the mussel holder (without the custom-made nylon mesh) was placed in the water tank (not shown in Fig. 5). The mussel holder was a 6.2 × 6.2 cm large frame to hold the mussel shell in position. The focus point, which was defined as the position of maximum pressure, was found at a distance of 45 ± 1 mm to the applicator, which was in line with the experimental setup of exposing mussels to ESWs (c.f. Fig. 2).

**Fig. 5.**
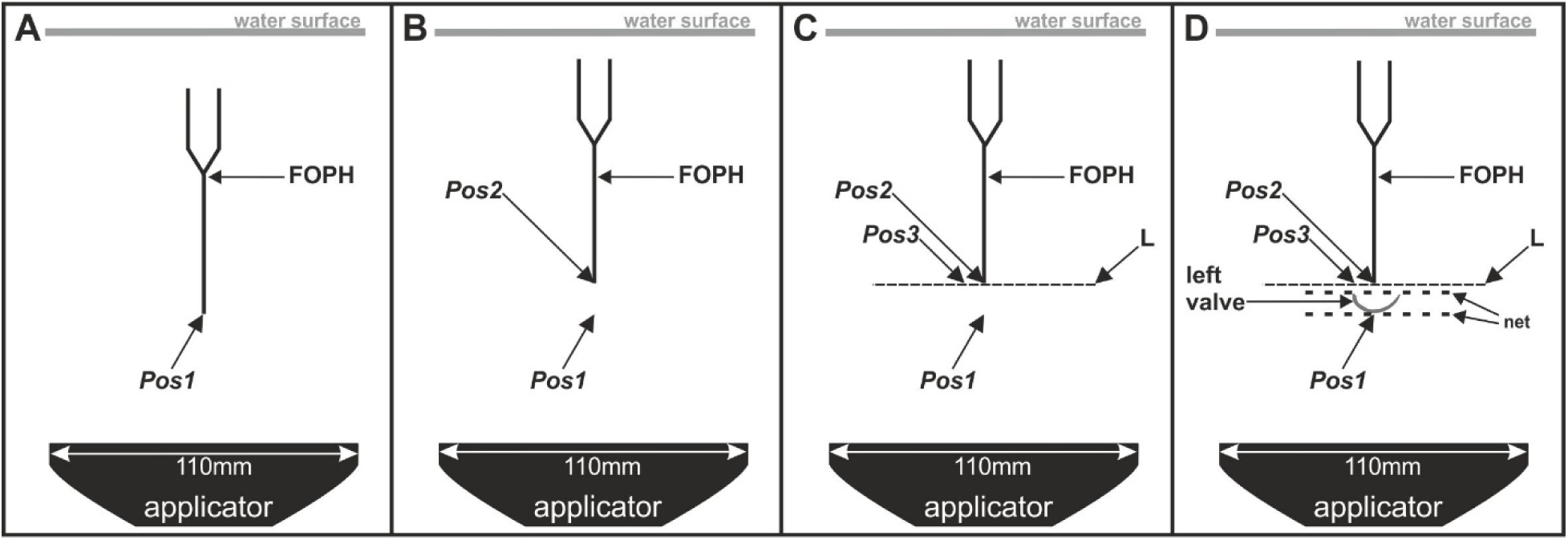
Scheme of the experimental setup of the acoustic measurements performed in the present study. The panels show the setup of the first (A), second (B), third (C) and fourth (D) series of performed acoustic measurements. Details are provided in the main text.

A second series of acoustic measurements (depicted in Fig. 5B) was dedicated to determining the pressure and ED at a height of 9.5 mm above the focus point within the 3D pressure field generated by the ESWs (indicated as “Pos2” in Fig. 5) (note that Pos2 approximately corresponded to the position of the middle of the right valve of a mussel in case the entire mussel shell were positioned in the 3D pressure field). To this end, the water tank was filled with tap water (mean temperature during the measurements: 11.5 ± 1.0 °C; water level 14.5 cm above the baseplate of the water tank).

A third series of acoustic measurements (depicted in Fig. 5C) was dedicated to determining the pressure and ED along a line (indicated as “L” in Fig. 5) that crossed Pos2 and was perpendicular to the acoustic axis (note that this line fully characterized the rotationally symmetrical 3D pressure field generated by the ESWs at a height of 9.5 mm above the focus point). To this end the FOPH was moved orthogonally to the acoustic axis. A special point on this line, whose position was 6 mm lateral to the XY position of the focus point, is indicated as “Pos3” in Fig. 5. The conditions in the water tank were the same as described above for the second series of acoustic measurements.

A fourth series of acoustic measurements was dedicated to determining the pressure and ED along Line “L” after placing the left valve of a zebra mussel in the pressure field as shown in Fig. 5D (i.e., with the outside of the valve facing the ESW applicator). The midpoint of the muscle valve was placed at Pos1, and the long axis of the valve was parallel to Line L (but at a different Z position). To ensure undisturbed sound propagation, attention was paid that no air bubbles were adherent to the mussel and the mussel holder. Again, the conditions in the water tank were the same as described above for the second series of acoustic measurements.

All measurements were repeated five times. The results were averaged and all non-focus signals were filtered with a low-pass filter at 5 Megahertz (MHz) while the focus signals were filtered with 100 MHz. Thereby deviations from the original data of <3 Megapascal (MPa) were attained, which corresponded to the signal noise.

### Statistical analysis

Mean and standard error of the mean (SEM) of all investigated variables (number of pixels in the linear pixel plots representing the hypostracum, average fluorescence signal intensity over the hypostracum, and results of the acoustic measurements) were calculated.

Differences in the mean number of pixels in the linear pixel plots representing the hypostracum between the left and the right valves as well as between positions 1 to 4 (c.f. Fig. 3C) were tested with two-way repeated measures ANOVA, with values obtained on the left and the right valve of each mussel at a given position 1 to 4 as matched data.

Differences in the average fluorescence signal intensity between mussels exposed to ESWs and mussels that were sham-exposed, as well as between the left and the right valves and between positions 1 to 4, were tested using a generalized linear model:

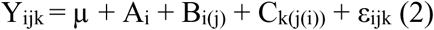

where µ was the mean and ε was the random error term. The fluorescence signal intensity (Y) was the dependent variable, the exposure to ESWs or sham exposure (A) and the side (B; left or right valve) were fixed effects. Mussels (C) were defined as random effects.

Spearman rank correlations were calculated in order to compare the mean fluorescence signal intensity within the left and the right valves at the different positions indicated in Fig. 3C. This was done for the left and the right valves with pooled data (1l/1r, 2l/2r, 3l/3r and 4l/4r) as well as for the left valves (1l, 2l, 3l and 4l) and the right valves (1r, 2r, 3u and 4r) separately.

The general linear model was calculated with the statistical software SAS (version 9.4; SAS Institute Inc., Cary, NC, USA). Spearman rank correlations and two way repeated measures ANOVA were calculated using IBM SPSS Statistics for Windows (Version 22.0; IBM, Armonk, NY, USA). P values smaller than 0.05 were considered statistically significant.

### Photography

Figure 1A (which is the basis for Fig. 3A) and Figure 3B were taken with a digital camera (Power Shot G12, Canon, Tokyo; Japan). The generation of Fig. 1C is described in detail above (BX51WI microscope; UPLSAPO20X objective; Olympus), as well as the generation of Fig. 3D and all panels in Figs 4, 7 and 8 (same microscope; UPLSAPO4X objective; Olympus). Figure 1D was taken with the same microscope using an UPLSAPO60XO objective (60×, oil, N.A. = 1.35) (Olympus). The final figures were constructed using Corel Photo-Paint X7 and Corel Draw X7 (both versions 17.5.0.907; Corel, Ottawa, Canada). Only minor adjustments of contrast and brightness were made using Corel Photo-Paint, without altering the appearance of the original materials.

## Results

### Thickness of the hypostracum

The mean thickness of the hypostracum (represented by mean numbers of pixels in the linear pixel plots found over the hypostracum) of all investigated mussels in Groups A and B significantly decreased from positions 1 (next to the umbo) to positions 4 (next to the shell growth zone) (1l: 89 ± 8; 2l: 83 ± 7; 3l: 76 ± 5; 4l: 56 ± 7; 1r: 116 ± 13; 2r: 77 ± 6; 3r: 67 ± 5; 4r: 58 ± 4; mean ± SEM) (p<0.001).

### Linear pixel plot analysis of fluorescence signal intensity

Results of the linear pixel plot analysis of calcein fluorescence imaging are summarized in Fig. 6; representative photomicrographs of calcein fluorescence imaging are shown in Fig. 7. Exposure to ESWs resulted in averaged 3.9-fold higher fluorescence signal intensity than sham exposure (based on averaging all measurement sites shown in Fig. 3C of a given mussel, resulting in a single value per mussel). This difference was statistically significant (p<0.001). No statistically significant difference was found in the mean fluorescence signal intensity between the left and the right valves of the mussels (p>0.67) despite the fact that the left valves were exposed to an approximately ten times higher energy density of the ESWs than the right valves. Besides this, after exposure to ESWs higher mean signal intensities were detected at the measurement sites 1l, 1r, 2l and 2r compared to the measurement sites 3l, 3r, 4l and 4r (Fig. 6). A statistically significant correlation was found between the fluorescence signal intensity and the measurement site considering both the left and the right valves (Spearman-Rho = −0.35; p<0.001). Considering only the right valves, the correlation was even higher (Spearman-Rho = −0.48; p<0.001). In contrast, no statistically significant correlation was found on the left site (Spearman-Rho = −0.24; p>0.05).

**Fig. 6.**
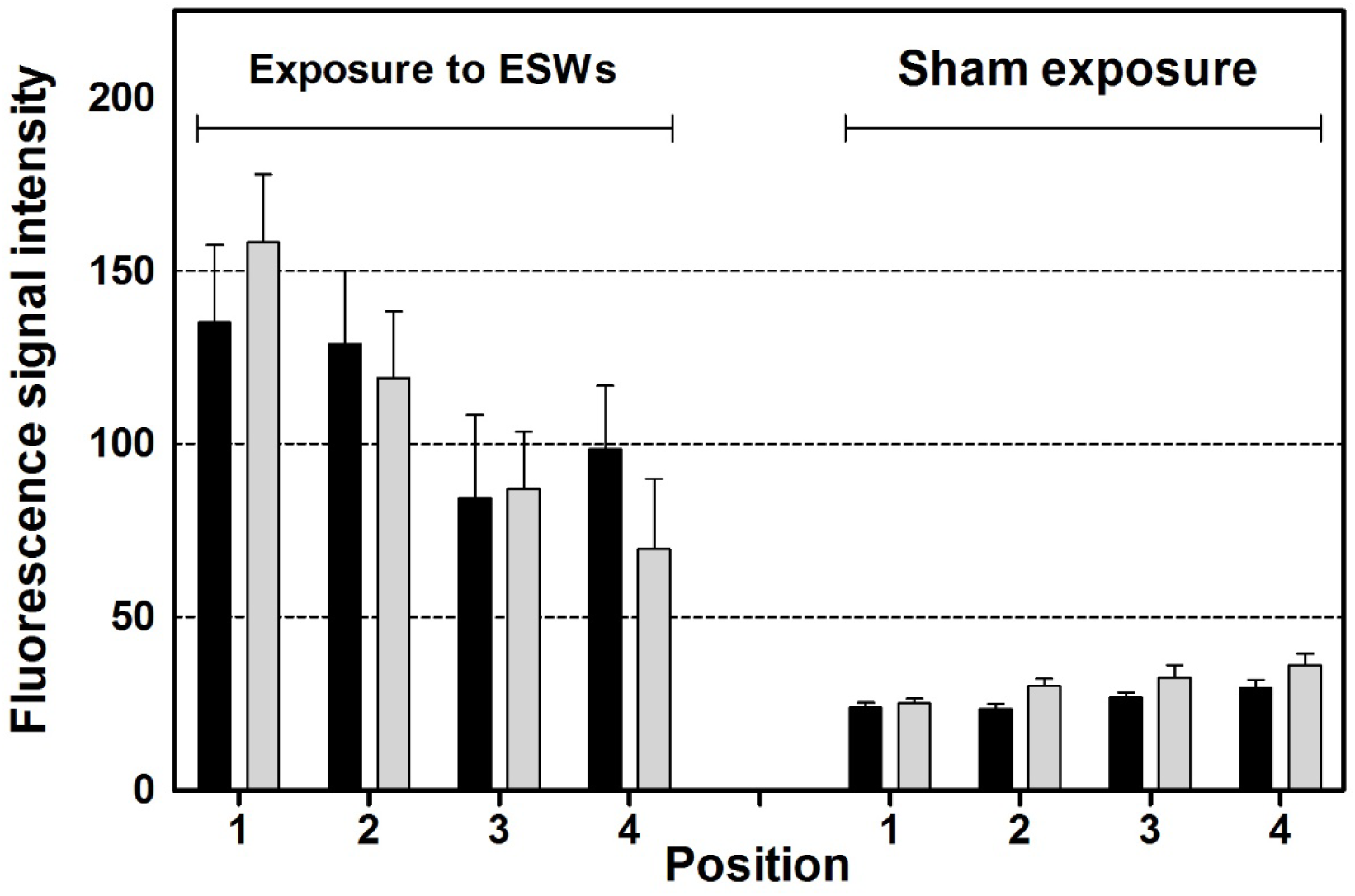
Results of quantitative analysis. The panel shows mean and standard error of the mean of the averaged fluorescence signal intensity (calcein fluorescence imaging) per pixel found at different positions (1-4; explained in detail in Fig. 3C) over the hypostracum of the left valves (closed bars) and the right valves (light gray bars) of the mussels two weeks after exposure to extracorporeal shock waves (ESWs) (on the left) or sham exposure (on the right) and incubation in calcein for 24 h immediately after exposure to ESWs or sham exposure.

**Fig. 7.**
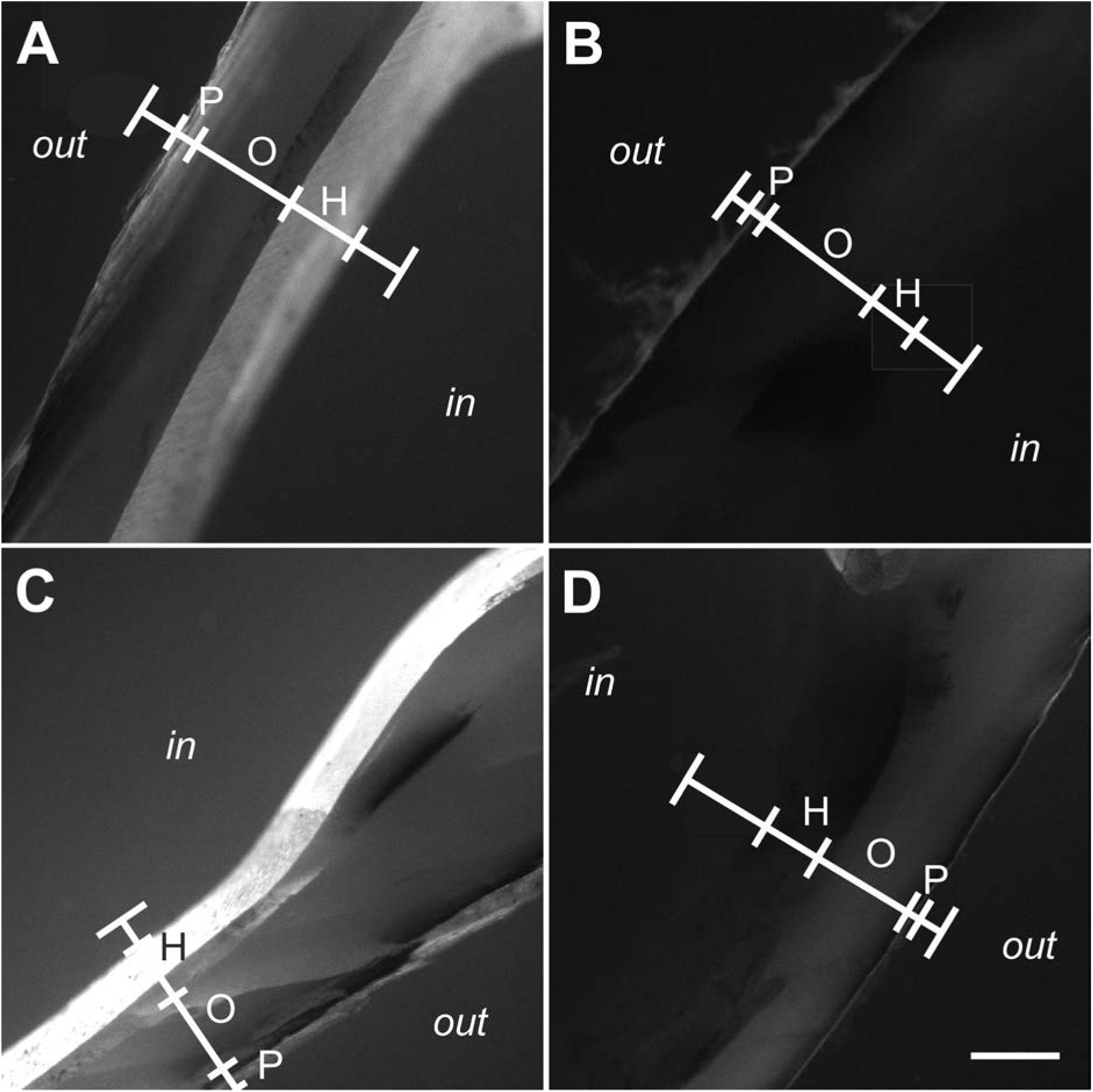
Fluorescence signal from calcein fluorescence imaging found in the shell of zebra mussels after exposure to extracorporeal shock waves. The panels show representative photomicrographs (calcein fluorescence imaging) of sections of the left (A,B) and right (C,D) valve of zebra mussels that were exposed to extracorporeal shock waves (A,C) or sham exposed (B,D). Photomicrographs were taken at positions 1l (A,B) and 1r (C,D) shown in Fig. 3C. The positions of linear pixel plot analysis (principle shown in Fig. 3D-F) are indicated (red lines). Abbreviations: out, outside surface of the mussel; P, periostracum; O, ostracum; H, hypostracum; in, inside surface of the mussel shell. The scale bar in (D) represents 250 µm in (A-D).

Representative photomicrographs of xylenol orange fluorescence imaging are shown in Fig. 8. Compared to calcein fluorescence imaging, the signal intensity found in xylenol orange fluorescence imaging was very weak and could not be quantified when imaging was performed with the camera calibrations established for calcein fluorescence imaging.

**Fig. 8.**
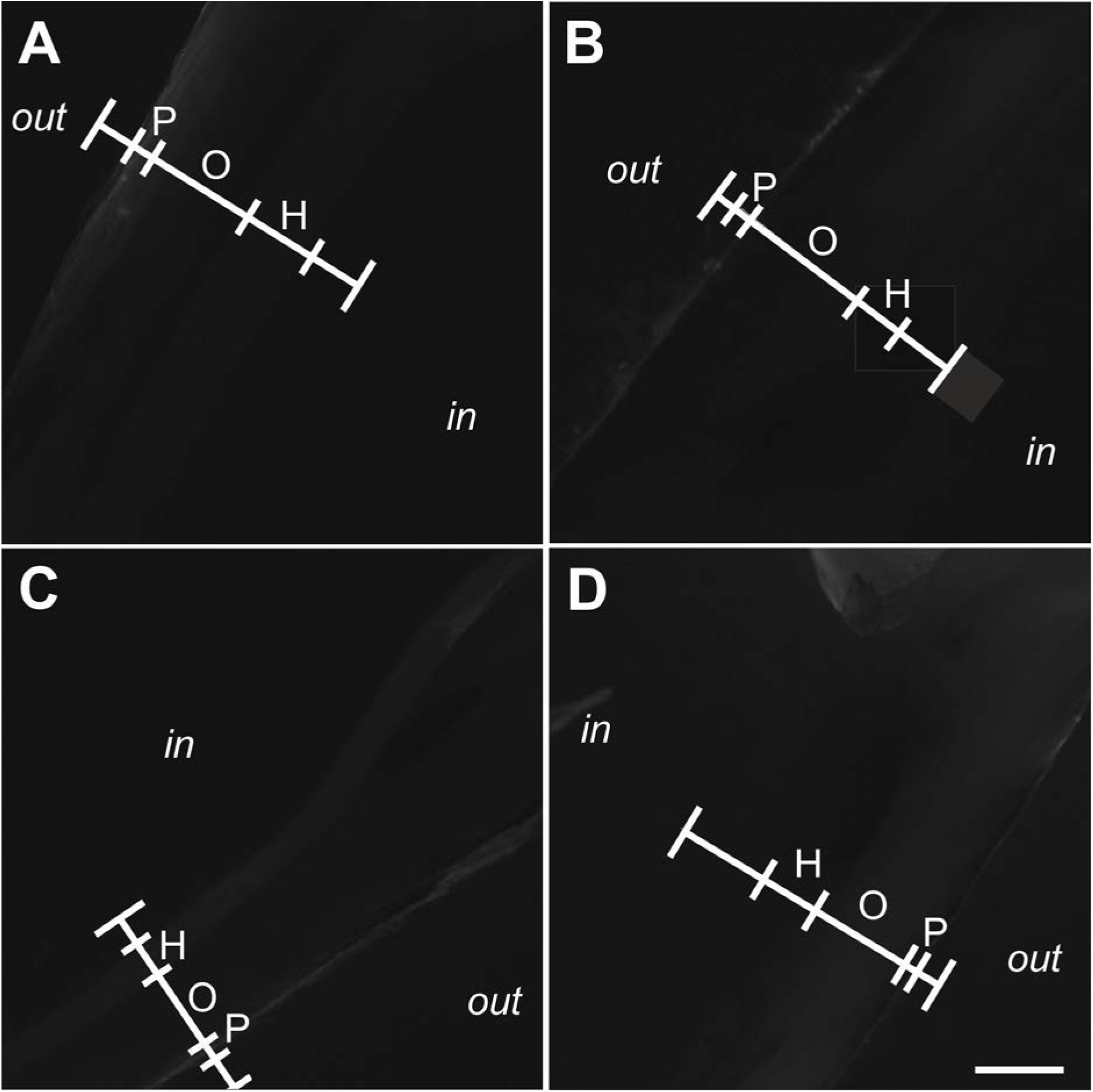
Absence of fluorescence signal from xylenol orange fluorescence imaging found in the shell of zebra mussels after exposure to extracorporeal shock waves. Representative photomicrographs (xylenol orange fluorescence imaging) of sections of the left (A,B) and right (C,D) valve of zebra mussels that were exposed to extracorporeal shock waves (A,C) or sham exposed (B,D). Photomicrographs were taken at positions 1l (A,B) and 1r (C,D) shown in Fig. 3C. The positions of linear pixel plot analysis (principle shown in Fig. 3D-F) are indicated (red lines). Abbreviations: out, outside surface of the mussel; P, periostracum; O, ostracum; H, hypostracum; in, inside surface of the mussel shell. The scale bar in (D) represents 250 µm in (A-D).

### Detection of microcracks

All investigated valves showed some micro-cracks, with no correlation between the position of the micro-cracks and the signal intensity found in calcein fluorescence imaging. Accordingly, the micro-cracks had to be attributed to their general occurrence in mussel shells, or the handling of the valves during histologic processing rather than to exposure to ESWs or sham exposure.

### Acoustic measurements

In the first series of acoustic measurements it was found that at the focus point of the 3D pressure field of the ESWs (i.e., at Pos1), the maximum pressure P_+_ was approximately 110 MPa and the minimum pressure P_−_ was approximately −20 MPa (Fig. 9A). The second and third series of acoustic measurements showed that 9.5 mm above the focus point (i.e., at Pos2), P_+_ and ED_+_ were reduced by more than 90%, and P_−_ and ED_−_ by approximately 60%, compared to the pressure and energy density at Pos1 (Figs 9B and 10A,B). Placing the muscle valve in the pressure field as depicted in Fig. 5D further reduced P_+_ and P_−_ at Pos2 (Figs 9D,E and 10A,B), but to a lesser extent than by moving from Pos1 to Pos2. At Pos3, P_+_ and P_−_ were further reduced compared to Pos2, and ED_+_ and ED_−_ were so low at Pos3 that they could not be measured precisely (Fig. 10C, D).

**Fig. 9.**
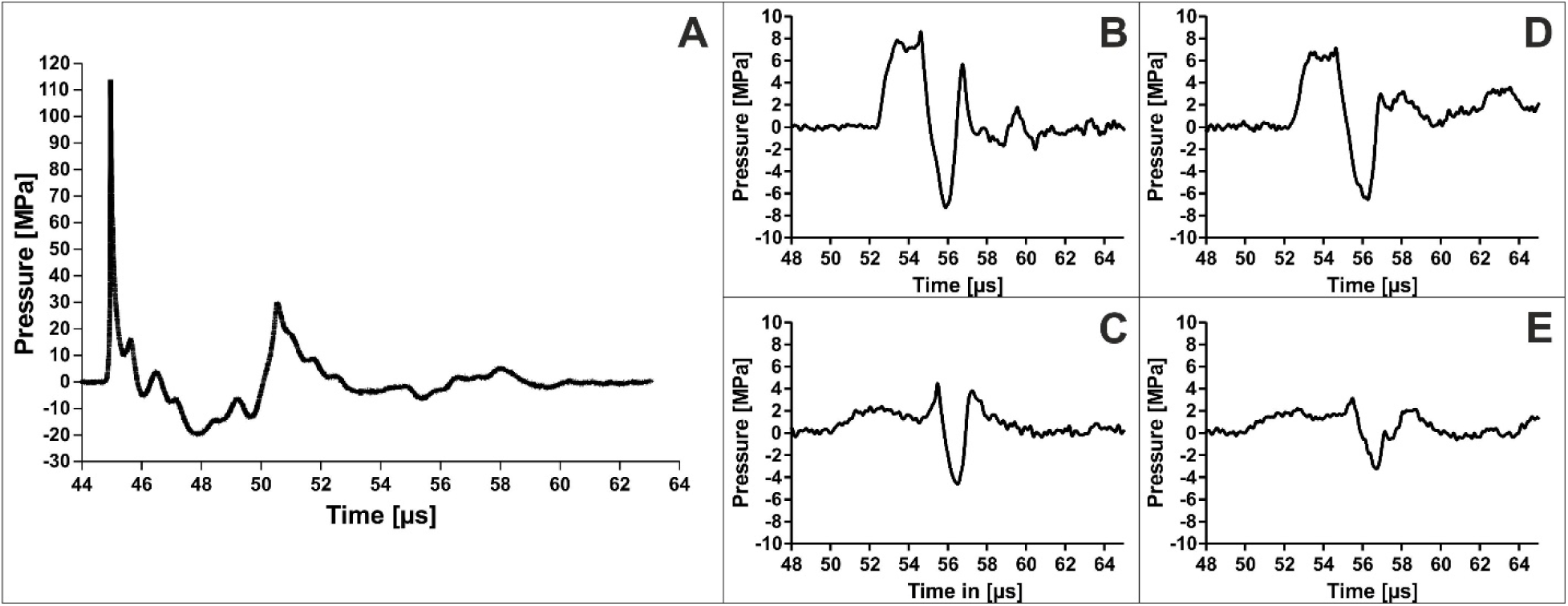
Results of acoustic measurements performed in the present study. The panels show pressure as a function of time of representative measurements of the three-dimensional pressure field of the investigated extracorporeal shock waves. (A) Pressure as a function of time at Pos1 during the first series of acoustic measurements (c.f. Fig. 5A). (B) Pressure as a function of time at Pos2 during the second series of acoustic measurements (c.f. Fig. 5B). (C) Pressure as a function of time at Pos3 during the third series of acoustic measurements (c.f. Fig. 5C). (D) Pressure as a function of time at Pos2 during the fourth series of acoustic measurements (c.f. Fig. 5D). (E) Pressure as a function of time at Pos3 during the fourth series of acoustic measurements (c.f. Fig. 5D).

**Fig. 10.**
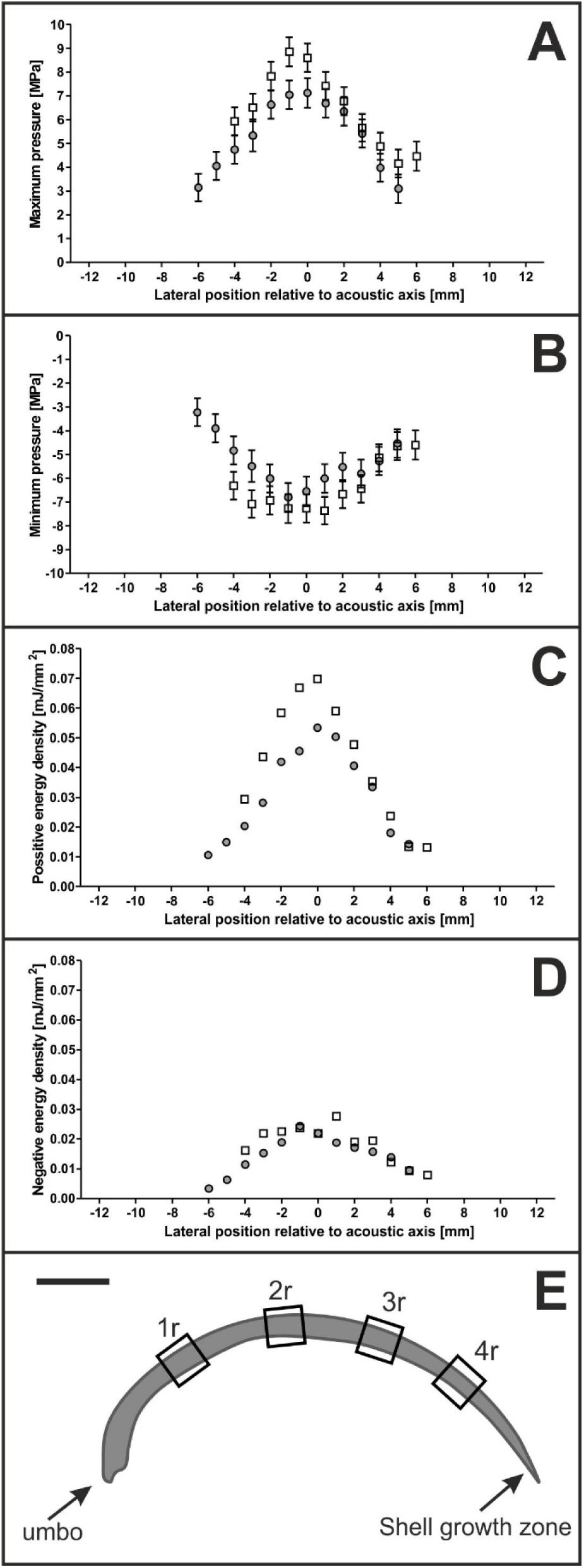
Results of acoustic measurements. The panels show mean and standard deviation of the maximum pressure P_+_ (A), minimum pressure P_−_ (B), positive energy density ED_+_ (C) and negative energy density ED_−_ (D) along Line L during the third (open squares) and fourth (dots) series of acoustic measurements (c.f. Fig. 5C,D). (E) shows a sketch of a section of a right mussle valve (drawn to scale), with the sites indicated where formation of new mineralized tissue after exposure to ESWs or sham exposure was investigated.

## Discussion

The results of this study allow several new insights into the complex process of induction of biomineralization by ESWs. First, exposure of zebra mussels to ESWs resulted in significantly increased incorporation of calcein and hence of increased shell reorganization or apposition within the hypostracum. Second, even a 10-fold difference in pressure exposure did not make a difference in terms of calcification comparing the left with the right valve of those mussels that were exposed to ESWs. Third, differences of the hypostracum thickness were correlated to the regional pattern of the measured reaction of the hypostracum across the valves (highest at positions next to the umbo and lowest at positions next to the shell growth zone) but were not correlated to the regional (3D) pressure distribution of the applied ESWs. This has significant clinical relevance as biomineralization is an important component of fracture healing.

Further, with regard to ESWT for fracture nonunions the present study supports the hypothesis that the biological reaction of the calcified tissue is not restricted to the position of the focus point of the ESWs. Although exact positioning of the focus point at the position of the fracture line is generally recommended (e.g. Furia et al., 2010; Schaden et al., 2001), experimental studies (Tischer et al., 2002; 2008) and clinical experience have shown that new bone formation can also occur outside of the focus zone. Treatment of fracture nonunions of superficial bones with radial ESWs (without focus point) yielded similar results (Kertzman et al., 2017; Silk et al., 2012). For the provider, ESWs produce an effect on bone more akin to a “shot gun” rather than a “rifle”.

### Formation of new mineralized tissue in mussels as a result of exposure to extracorporeal shock waves

The focus of the ESWs was small compared to the size of the mussels and, as a result, the pressure during exposure to ESWs showed substantial regional differences across the mussel shell (Figs 9,10). According to the manufacturer of the ESWT device used in this study the diameter of the 5 MPa focus of the ESWs is 20.8 mm in XY directions when operating this device at highest settings (i.e., ED = 0.4 mJ/mm^2^) as performed in this study. Our own acoustic measurements showed that 9.5 mm above the focus point (where the right valve would have been) the 5 MPa focus of the ESWs had a diameter of 10 mm in XY directions (Fig. 10A). Thus, the umbo of both the left and right valves of the mussels exposed to ESWs was always outside the 5 MPa focus. Furthermore, the umbo of both the left and right valves of the mussels exposed to ESWs was always outside the -6dB focus (diameter in XY direction: 2.4 mm; length in Z direction: 9.6 mm when operating the used ESWT device at ED = 0.4 mJ/mm^2^ according to the manufacturer of this ESWT device). This was also confirmed by our acoustic measurements since 9.5 mm above the focus point only 10% of P_+_ was measured. However, we found the highest calcein fluorescence signal intensity over the hypostracum in regions next to the umbo.

Because of their calcium binding characteristics, calcein and xylenol orange are commonly used markers in studies on vertebrate bone remodeling or bone growth *in vivo* (Rahn and Perren, 1971; Suzuki and Mathews, 1966; van Gaalen et al., 2010). In vertebrates, fluorescent dyes such as calcein were used to label bone and thus apposition of new cortical bone after exposure to ESWs (Delius et al., 1995). Calcein marking has also been proven to be a suitable tool in ecological and toxicological studies on mussels since the apposition of new shell material can easily be measured. For example, van der Geest et al. (2011) exposed mussels to calcein in order to measure the distance from the growing edge at the time of calcein exposure (marked by a calcein band) to the growing edge three months later.

In this study, calcein was used for the first time as *in vivo* marker for mussel shell modification of the hypostracum after physical disturbance, i.e., exposure to ESWs. We could show increased calcein fluorescence signal intensity not only in a small calcein band built next to the shell growth zone (Fig. 3B) but across the whole hypostracum, i.e. from the umbo to the growth zone (Figs 3D and 6). We also quantified the fluorescence signal intensity to show differences in the amount of processed calcium within the hypostracum after exposure to ESWs.

### Characteristics of the pressure field generated by ESWs versus spatial distribution of the biological response across the hypostracum

Unexpectedly, the regional pattern of calcein fluorescence signal intensity correlated with the thickness of the mussel shell but not with the spatial distribution of the pressure and energy density of the applied ESWs. The measured calcein fluorescence signal intensity was highest next to the umbo. The hypostracum of the umbo has more nacreous layers, i.e. is thicker, than the hypostracum next to the growth zone, because a characteristic of the freshwater mussel hypostracum is that new nacreous layers are added during growth lateral of the extrapallial space (e.g., Geist et al., 2005; Immel et al., 2016; Lindh et al., 1988). The higher number of pixels next to the umbo compared to the decreasing number of pixels towards the shell growth zone found in this study is in line with the expected shell thickness caused by the natural growth of mussel shells.

Assuming that mussels perform locally restricted biocalcification, e.g. after shell damage (Beedham, 1964; Mount et al., 2004), ESWs may trigger a different mechanism that results in a biological response that is not correlated with the spatial distribution of the energy density of the applied ESWs. Even though the maximum pressure at the left valve (i.e., at the focus point) was approximately ten times higher than at the right valve, there was no difference in the mean fluorescence signal intensities between the left and the right valves. It is unlikely that even very low shock wave energy (e.g., P_+_ < 2 MPa at position 1r; c.f. Fig. 10) is sufficient to stimulate formation of new mineralized tissue locally. However, wave reflections at the inner right mussel valve might increase the pressure on the left valve during the *in vivo* experiments. Although this effect could not be measured during the acoustic measurements as only the left valve could be used, we do not expect a major influence on the pressure field. Rather, it is more likely that the mechanical stimulus generated at the focus point was physically transferred through the mussel shell and/or activated a biological response that was transmitted through the mussel soft tissue. This mechanism could be a mechanical wave that is propagated from the focus point through the shell itself, activating cells directly next to the shell.

A mechanical impact next to the focus point resulting in a biological response not limited to the position of the focus point but physiologically distributed across the mussel soft tissue could explain the observed regional pattern of calcein fluorescence signal intensity after exposure to ESWs. Cells in the mantle of the mussel (sensors) could be activated by a biological signal after exposure to ESWs. The conversion of the acoustic energy of ESWs into a biological signal was suggested to be a result of the mechanotransduction in the soft tissue after stimulation of hard tissue (Wang et al., 2003). Afterwards, several ways of information transfer from sensor to effector are conceivable to initiate a reaction across the whole mussel shell.

The information could be passed through the organism by the nervous system to the effector cells in the mantel epithelium. For vertebrate bones it is well known that sensory and sympathetic nerve fibers are critically involved in bone development, growth and remodeling (e.g., Chenu, 2004). The mantle epithelium cells and the granular hemocytes regulate the biocalcification in mussels. The outer mantle epithelium of the mussel mantle regulates calcification during growth within the extrapallial space by producing organic material and controlling the transport of Ca_2_^+^ as well as further required ions to the extrapallial space (e.g. Beedham 1964, Immel et al. 2016). After shell injuries granular hemocytes (i.e., amoeboid cells with macrophage-like functions) accumulate at the damaged spots and support shell regeneration (e.g. Kádár, 2008; Mount et al., 2004). The mechanisms how the granular hemocytes are directed to the shell are not fully understood and could be initiated by nerves or by transmitters that circulate within the haemolymphe.

### Benefits of zebra mussels as novel animal model in basic research on ESWT

So far, research into the molecular and cellular mechanisms of new bone formation and biocalcification after exposure of bone to ESWs has to the best knowledge only been conducted in vertebrate models (e.g. Bulut et al., 2006; Rompe et al., 2001; van der Jagt et al., 2011). Tischer et al. (2002) demonstrated that ESWs can induce new bone formation inside and outside the focus zone. In fact, the important question about correlations between the regional (3D) distribution of pressure generated by ESWs and the regional (3D) pattern of new bone formation induced by ESWs could not be answered. This is due to the fact that for technical reasons, it is not possible to measure the regional distribution of pressure generated by ESWs within a limb of a human or one of the commonly used animal models in ESWT research (goat, rabbit, rat and mouse) *in vivo*.

In search of a novel animal model that could provide answers to the question about correlations between the regional (3D) distribution of pressure generated by ESWs and the regional (3D) pattern of new bone formation induced by ESWs it is important to note that mineralization in biological systems is a genetically and physiologically regulated process (Addadi and Weiner, 1992; Geist et al., 2005). In vertebrate bones, mechanosensitive osteocytes detect mechanical signals and consequently stimulate osteoblasts to initiate new bone formation and biocalcification. This mechanism is not only activated in directly impaired parts, but also in not directly affected parts of the bone (Klein-Nulend et al., 2013).

However, studies on vertebrate bones are hampered by the organization of the vertebrate bone itself. The mechanosensors (i.e., the osteocytes) are located within the mineralized bone matrix, limiting accessability. The extraction of living osteocytes followed by *in vitro* studies on the biological response of the mechanosensors and their complex interplay with the effectors (i.e., the osteoblasts and osteoclasts) in vertebrate bone after exposure to certain stimuli such as ESWs is therefore considered infeasible (Klein-Nulend et al., 2013).

Furthermore, studies on the effects of ESWs were so far exclusively done on the effectors (osteoblasts) of the vertebrate bone. A direct impact of ESWs on the osteoblasts was only shown *in vitro* (e.g. Hofmann et al., 2008; Wang et al., 2001). Access to the mechanosensors (osteocytes) is strictly limited to the *in vivo* situation. It has been suggested that altered morphology of the vertebrate bone mechanosensors results in bone remodeling or induction of bone repair mechanisms (e.g. Bulut et al., 2006; Burr, 2002). For instance, osteocytes sense the mechanical load of shear forces and transform it into biological signals (Klein-Nulend et al., 2013). A suggested physical effect of ESWs on biological tissue are shear forces (e.g. Delacretaz et al., 1995), and it appears crucial to study ESW-induced shear stress affecting the sensor cells. The mechanical forces produced by ESWs can result in extreme pressure fluctuations and, under specific conditions, even in the formation of cavitation that can generate so-called “jet streams”, which are high velocity liquid streams (Delacrétaz et al., 1995; Gerdesmeyer et al., 2002, Ueberle, 2016). Such jet streams can produce substantial shear forces, which are suggested to activate osteocytes and thus new bone formation (e.g. Klein-Nulend et al., 2013).

The principles of biocalcification in invertebrates with calcified tissues, particularly mussels, show despite their different mineral types many similarities to those observed in vertebrate bone (e.g., Addadi et al., 2006; Mahamid et al., 2011; Immel et al., 2016). In fact, mussels appear to be an attractive novel animal model for studying the impact of ESWs on biocalcification because, unlike in vertebrates, the mechanosensors and effectors are not embedded within the calcified tissue of the skeleton (i.e., the mussel shell). Instead, the relevant cells (i.e., the outer mantle epithelium cells) are located in the mantle tissue of the mussel or originate from the mantle (Fig. 1). These cells are crucial for the production of new shell material (e.g., Immel et al., 2016; Machado et al., 1988a). Between the mantle and the shell is an extrapallial space, where the mineralization takes place. This space shows a striking similarity to the mineralization zone in bone of vertebrates. The biocalcification of the inner shell layer (hypostracum) is controlled by epithelium cells of the mantle. These cells are sensitive to physical as well as chemical stimuli (being a sensor) and secrete nacre to the hypostracum (being an effector), which was previously observed under laboratory conditions (Lopes-Lima et al., 2008; Machado et al., 1988b; Soares-da-Silva et al., 1998). The biocalcification performed by the mantle epithelium cells is similar to the action of osteoblasts in the cambium layer of the periosteum or the odontoblasts building dentin. Analog to human bone, the mussel shell is characterized as calcified tissue yet without living cells and with aragonite and not hydroxyl apatite as mineralized matrix (Pathy and Mackie, 1993).

Hence, the most important advantage of using mussels for research into biocalcification is the access to sensor and effector cells that perform the biological response of calcifying tissue to ESWs. These characteristics render *Dreissena polymorpha* a very attractive model for studying biological effects of ESWs. For example, a long standing question is whether ESWs with identical energy density at the focus point but generated with respectively electrohydraulic, electromagnetic or piezoelectric ESWT devices (summarized in, e.g., Schmitz et al., 2015) exert the same biologic responses on calcifying tissue. Another question is whether a low number of ESWs with high positive energy density exerts the same biologic responses on calcifying tissue than a high number of ESWs with low energy density. Furthermore, *Dreissena polymorpha* can be used to investigate effects of ESWs in combination with the administration of drugs and biological agents.

## Acknowledgments

We thank Verena Huber, Katharina Stoeckl, Jörg Steinhilber, Claudia Harbauer, Barbara Mosler, Bernd Riedelsheimer and Beate Aschauer for technical support.

## References

Addadi, L. and Weiner, S. (1992). Control and design principles in biological mineralization. Angew. Chem. Int. 31, 153–169.

Addadi, L., Joester, D., Nudelman, F. and Weiner, S. (2006) Mollusk shell formation: a source of new concepts for understanding biomineralization processes. Chemistry. 12, 980–987.

Beedham, G. (1964) Repair of the shell in species of Anodonta. Proc. Zool. Soc. Lond. 145, 107–123.

Beggel, S., Cerwenka, A.F., Brandner, J. and Geist J. (2015). Shell morphological versus genetic identification of quagga mussel (*Dreissena bugensis*) and zebra mussel (*Dreissena polymorpha*). Aquat. Invasions 10, 93–99.

Bulut, O., Eroglu, M., Ozturk, H. and Tezeren, G. (2006). Extracorporeal shock wave treatment for defective nonunion of the radius: a rabbit model. J. Orthop. Surg. 14, 133–137.

Burr, D.B. (2002). Targeted and nontargeted remodeling. Bone 30, 2–4

Chamberlain, G.A. and Colborne, G.R. (2016). A review of the cellular and molecular effects of extracorporeal shockwave therapy. Vet. Comp. Orthop. Traumatol. 29, 99–107.

Checa, A. (2000). A new model for periostracum and shell formation in Unionidae (Bivalvia, Mollusca). Tissue Cell 32, 405–416.

Cheng, J.H. and Wang, C.J. (2015). Biological mechanism of shockwave in bone. Int. J. Surg. 24, 143–146.

Chenu, C. (2004). Role of innervation in the control of bone remodeling. J. Musculoskelet. Neuronal. Interact. 4, 132–134.

Claxton WT, Martel A, Dermott RM, Boulding EG. (1997). Discrimination of field-collected juveniles of two introduced dreissenids (*Dreissena polymorpha* and *Dreissena bugensis*) using mitochondrial DNA and shell morphology. Can. J. Fish. Aquat. Sci. 54, 1280–1288.

Da Costa Gómez, T.M., Radtke, C.L., Kalscheur, V.L., Swain, C.A., Scollay, M.C., Edwards, R.B., Santschi, E.M., Markel, M.D. and Muir, P. (2004). Effect of focused and radial extracorporeal shock wave therapy on equine bone microdamage. Vet. Surg. 33, 49–55.

Delacrétaz, G., Rink, K., Pittomvils, G., Lafaut, J.P., Vandeursen, H. and Boving, R. (1995). Importance of the implosion of ESWL-induced cavitation bubbles. Ultrasound Med. Biol. 21, 97–103.

Delius, M., Draenert, K., Al Diek, Y. and Draenert, Y. (1995). Biological effects of shock waves: In vivo effect of high energy pulses on rabbit bone. Ultrasound Med. Biol. 95; 21,1219–25.

Dietz-Laursonn, K., Beckmann, R., Ginter, S., Radermacher, K., and de la Fuente, M. (2016). In-vitro cell treatment with focused shockwaves-influence of the experimental setup on the sound field and biological reaction. J. Ther. Ultrasound 4, 1–14.

Furia, J.P., Juliano, P.J., Wade, A.M., Schaden, W., Mittermayr, R. (2010). Shock wave therapy compared with intramedullary screw fixation for nonunion of proximal fifth metatarsal metaphyseal-diaphyseal fractures. J. Bone Joint Surg. Am. 92, 846–854.

Geist, J., Auerswald, K. and Boom, A. (2005). Stable carbon isotopes in freshwater mussel shells: Environmental record or marker for metabolic activity? Geochim. Cosmochim. Acta. 69, 3545–3554.

Gerdesmeyer, L., Maier, M., Haake, M. and Schmitz C. (2002). Physikalisch-technische Grundlagen der extrakorporalen Stosswellentherapie. Orthopade 31, 610–617.

Gollwitzer, H., Gloeck, T., Roessner, M., Langer, R., Horn, C., Gerdesmeyer, L. and Diehl, P. (2013). Radial extracorporeal shock wave therapy (rESWT) induces new bone formation *in vivo*: results of an animal study in rabbits. Ultrasound Med. Biol. 2013; 39, 126–133.

Haupt, G. (1997). Use of extracorporeal shock waves in the treatment of pseudarthrosis, tendinopathy and other orthopedic diseases. J. Urol. 158, 4–11.

Hofmann, A., Ritz, U., Hessmann, M.H., Alini, M., Rommens, P.M., Rompe, J.-D. (2008). Extracorporeal shock wave-mediated changes in proliferation, differentiation, and gene expression of human osteoblasts. J. Trauma 65, 1402–1410.

Ikeda, K., Tomita, K., Takayama, K. (1999). Application of extracorporeal shock wave on bone: preliminary report. J. Trauma 47, 946–950.

Immel, F., Broussard, C., Catherinet, B., Plasseraud, L., Alcaraz, G., Bundeleva, I. and Marin, F. (2016). The shell of the invasive bivalve species *Dreissena polymorpha*: biochemical, elemental and textural investigations. PLoS One 11, e0154264.

Ingersoll, C.G., Augspurger, T.P., Barnhart, C., Bidwell, J., Bishop, C., Black, M., Cope, G., Bringolf, R.B., Dwyer, F.J., Greer, I.E., Keller, A., Linder, G., Neves, R.J., Newton, T.J., Pepino, R., Roberts, A., Roberts, C., Salazar, M., Samuel, A., Stephan, C.D., Van Hassel, J.H. and Wang, N. (2006) Standard guide for conducting laboratory toxicity tests with freshwater mussels. Special Technical Report ASTM E2455-06. DOI:10.1520/E2455-06R13.

Jantz, B. and Neumann, D. (1998). Growth and reproductive cycle of the zebra mussel in the River Rhine as studied in a river bypass. Oecologia 114, 213–225.

Kádár, E. (2008) Haemocyte response associated with induction of shell regeneration in the deep-sea vent mussel Bathymodiolus azoricus (Bivalvia: Mytilidae). J. Exp. Mar. Bio. Ecol. 362,71–78.

Kertzman, P., Császár, N.B.M., Furia, J.P. and Schmitz, C. (2017) Radial extracorporeal shock wave therapy for the treatment of fracture nonunions: a case series. J. Orthop. Surg. Res. 12, 164.

Klein-Nulend, J., Bakker, A.D., Bacabac, R.G., Vatsa, A. and Weinbaum, S. (2013). Mechanosensation and transduction in osteocytes. Bone 54, 182–90.

Lindh, U., Mutvei, H., Sunde, T., and Westermark, T. (1988). Environmental history told by mussel shells. Nucl. Instrum. Methods Phys. Res. B. 30, 388–392.

Lopes-Lima, M., Bleher, R., Forg, T., Hafner, M. and Machado, J. (2008). Studies on a PMCA-like protein in the outer mantle epithelium of *Anodonta cygnea*: insights on calcium transcellular dynamics. J. Comp. Physiol. B. 178, 17–25.

Machado, J., Castilho, F., Coimbra, J., Monteiro, E., Sá, C. and Reis, M. (1988a) Ultrastructural and cytochemical studies in the mantle of *Anodonta cygnea*. Tissue Cell 20, 797–807.

Machado, J., Coimbra, J., Sá, C. and Cardoso, I. (1988b). Shell thickening in *Anodonta cygnea* by induced acidosis. Comp. Biochem. Phys. A. 91, 645–651.

Mahamid, J., Addadi, L. and Weiner, S. (2011). Crystallization pathways in bone. Cells Tissues Organs 194, 92–97.

Mann, S. (1988). Molecular recognition in biomineralization. Nature 332, 119–124.

Milz, S. and Putz, R. (1994). Quantitative morphology of the subchondral plate of the tibial plateau. J. Anat. 185, 103–110.

Mount, A.S., Wheeler, A.P., Paradkar, R.P. and Snider, D. (2004). Hemocyte-mediated shell mineralization in the Eastern Oyster. Science 304, 297–300.

O’Brien, F.J., Taylor, D. and Lee, T.C. (2002). An improved labelling technique for monitoring microcrack growth in compact bone. J. Biomech. 35, 523–526.

Pautke, C., Vogt, S., Tischer, T., Wexel, G., Deppe, H., Milz, S., Schieker, M. and Kolk, A. (2005). Polychrome labeling of bone with seven different fluorochromes: enhancing fluorochrome discrimination by spectral image analysis. Bone 37, 441–445.

Pathy, D.A., Mackie, G.L. (1993). Comparative shell morphology of *Dreissena polymorpha, Mytilopsis leucophaeata*, and the “quagga” mussel (Bivalvia: Dreissenidae) in North America. Can. J. Zool. 71, 1012–1023.

Petit, H., Davis, W.L., Jones, R.G. and Hagler, H.K. (1980). Morphological studies on the calcification process in the fresh-water mussel *Amblema*. Tissue Cell 12, 13–28.

Rahn, B.A. and Perren, S.M. (1971). Xylenol orange, a fluorochrome useful in polychrome sequential labeling of calcifying tissues. Stain. Technol. 46, 125–129.

Rompe, J.D., Rosendahl, T., Schöllner, C., Theis, C. (2001). High-energy extracorporeal shock wave treatment of nonunions. Clin. Orthop. Relat. Res. 387, 102–111.

Schaden, W., Fischer, A. and Sailler, A. (2001). Extracorporeal shock wave therapy of nonunion or delayed osseous union. Clin. Orthop. Relat. Res. 387, 90–4.

Schaden, W., Mittermayr, R., Haffner, N., Smolen, D., Gerdesmeyer, L., and Wang, C.J. (2015). Extracorporeal shockwave therapy (ESWT) – First choice treatment of fracture non-unions?. Int. J. Surg. 24, 179–183.

Schmitz, C., Császár, N.B., Rompe, J.D., Chaves, H., Furia, J.P. (2013). Treatment of chronic plantar fasciopathy with extracorporeal shock waves (review). J. Orthop. Surg. Res. 8, 31.

Schmitz, C., Császár, N.B., Milz, S., Schieker, M., Maffulli, N., Rompe, J.D. and Furia, J.P. (2015). Efficacy and safety of extracorporeal shock wave therapy for orthopedic conditions: a systematic review on studies listed in the PEDro database. Br. Med. Bull. 116, 115–138.

Silk, Z.M., Alhuwaila, R.S. and Calder, J.D. (2012). Low-energy extracorporeal shock wave therapy to treat lesser metatarsal fracture nonunion: case report. Foot Ankle Int. 33, 1128–1132.

Soares-da-Silva, I.M., Almeida, M.J., Serrão, P.M., Coelho, M.A. and Machado, J. (1998). l-3,4-dihydroxyphenylalanine (l-DOPA) in *Anodonta cygnea*: variation with acidosis. Comp. Biochem. Physiol. A. Mol. Integr. Physiol. 120, 463–468.

Suzuki, H.K. and Mathews, A. (1966). Two-color fluorescent labeling of mineralizing tissues with tetracycline and 2,4-bis[N,NV-di-(carbomethyl)aminomethyl] fluorescein. Stain. Technol. 41, 57–60.

Tischer, T., Milz, S., Anetzberger, H., Müller, P., Wirtz, D., Schmitz, C., Ueberle, F. and Maier, M. (2002). Extrakorporale Stosswellen induzieren ventral-periostale Knochenneubildung ausserhalb der Fokuszone - Ergebnisse einer in-vivo-Untersuchung am Tiermodell. Z. Orthop. Ihre. Grenzgeb. 140, 281–5.

Tischer, T., Milz, S., Weiler, C., Pautke, C., Hausdorf, J., Schmitz, C., and Maier, M. (2008). Dose-dependent new bone formation by extracorporeal shock wave application on the intact femur of rabbits. Eur. Surg. Res. 41, 44–53.

Ueberle, F. (2016) Einsatz von Stoßwellen in der Medizin. In Medizintechnik: Verfahren - Systeme - Informationsverarbeitung. (ed. R. Kramme), pp. 1–37. Berlin:Springer Verlag Berlin Heidelberg.

Valchanou, V.D. and Michailov, P. (1991). High energy shock waves in the treatment of delayed and nonunion of fractures. Int. Orthop. 15, 181–184.

van der Geest, M., van Gils, J.A., van der Meer, J., Olff, H. and Piersma, T. (2011). Suitability of calcein as an in situ growth marker in burrowing bivalves. J. Exp. Mar. Biol. Ecol. 399, 1–7.

van der Jagt, O.P., Piscaer, T.M., Schaden, W., Li, J., Kops, N., Jahr, H., van der Linden, J.C., Waarsing, J.H., Verhaar, J.A.N., de Jong, M. and Weinans, H. (2011). Unfocused extracorporeal shock waves induce anabolic effects in rat bone. J. Bone Joint Surg Am. 93, 38–48.

van Gaalen, S.M., Kruyt, M.C., Geuze, R.E., de Bruijn, J.D., Alblas, J. and Dhert, W.J. (2010). Use of fluorochrome labels in in vivo bone tissue engineering research. Tissue Eng. Part. B. Rev. 16, 209–217.

Wang, C-J., Chen, H-S., Chen, C-E. and Yang, K.D. (2001). Treatment of nonunions of long bone fractures with shock waves. Clin. Orthop. Relat. Res. 387, 95–101.

Wang, F.S., Yang, K.D., Kuo, Y.R., Wang, C.J., Sheen-Chen, S.M., Huang. H.C. and Chen, Y.J. (2003). Temporal and spatial expression of bone morphogenetic proteins in extracorporeal shock wave-promoted healing of segmental defect. Bone 32, 387–96.

Zelle, B.A., Gollwitzer, H., Zlowodzki, M. and Buhren, V. (2010). Extracorporeal shock wave therapy: current evidence. J. Orthop. Trauma 24, 66–70.

